# A layered, hybrid machine learning analytic workflow for mouse risk assessment behavior

**DOI:** 10.1101/2022.08.22.504822

**Authors:** Jinxin Wang, Paniz Karbasi, Liqiang Wang, Julian P. Meeks

## Abstract

Accurate and efficient quantification of animal behavior facilitates the understanding of the brain. An emerging approach within machine learning (ML) field is to combine multiple ML-based algorithms to quantify animal behavior. These so-called hybrid models have emerged because of limitations associated with supervised (e.g., random forest, RF) and unsupervised (e.g., hidden Markov model, HMM) ML classifiers. For example, RF models lack temporal information across video frames, and HMM latent states are often difficult to interpret. We sought to develop a hybrid model, and did so in the context of a study of mouse risk assessment behavior. We utilized DeepLabCut to estimate the positions of mouse body parts. Positional features were calculated using DeepLabCut outputs and were used to train RF and HMM models with equal number of states, separately. The per-frame predictions from RF and HMM models were then passed to a second HMM model layer (“reHMM”). The outputs of the reHMM layer showed improved interpretability over the initial HMM output. Finally, we combined predictions from RF and HMM models with selected positional features to train a third HMM model (“reHMM+”). This reHMM+ layered hybrid model unveiled distinctive temporal and human-interpretable behavioral patterns. We applied this workflow to investigate risk assessment to trimethylthiazoline and snake feces odor, finding unique behavioral patterns to each that were separable from attractive and neutral stimuli. We conclude that this layered, hybrid machine learning workflow represents a balanced approach for improving the depth and reliability of ML classifiers in chemosensory and other behavioral contexts.

**Significance Statement:** In this study, we integrate two widely-adopted machine learning (ML) classifiers, random forest and hidden Markov model, to develop a layered, hybrid ML-based workflow. Our workflow not only overcomes the intrinsic limitations of each model alone, but also improves the depth and reliability of ML models. Implementing this analytic workflow unveils distinctive and dynamic mouse behavioral patterns to chemosensory cues in the context of mouse risk assessment behavioral experiments. This study provides an efficient and interpretable analytic strategy for the quantification of animal behavior in diverse experimental settings.

## Introduction

Behavior is the muscular output of an organism reflecting the function of the central nervous system (CNS) (Schlinger, 2015). Accurately and efficiently quantifying animal behavior improves our knowledge of the structural and functional connectivity of the CNS underlying complex behaviors (Krakauer et al., 2017). In general, videography-based recording and manual annotation have long been a standard method for animal behavior quantification (Anderson and Perona, 2014). Recently, advancements in machine learning (ML) software have enabled tracking of animal/body parts of interest movement automatically in video recordings (Berman, 2018; Mathis et al., 2018b). Of these, DeepLabCut, built based on transfer learning with deep neural networks, has been widely adopted due to its efficient and intuitive framework, and supports markerless movement tracking and pose estimation (Mathis et al., 2018a; Nath et al., 2019).

Regardless of the specific approach used for tracking animal positions and poses, a major challenge remains for those seeking to quantify complex animal behaviors. Typically, the multidimensional matrix of feature positions produced by DeepLabCut or other tracking methods is used as the input for additional algorithmic tools that perform dimensionality reduction, feature recognition, and ultimately a prediction of an animal’s behavioral state at each point in space and time (Dell et al., 2014; Guo et al., 2015; Machado et al., 2015). One of these tools, the Random Forest (RF), a versatile supervised ML algorithm, has been used in the context of human (Siddiqui and Medioni, 2010; Trinh et al., 2012) and mouse (Hong et al., 2015; Winters et al., 2022) behavioral analysis. RFs generally have high classification accuracy, a capacity to fit complex datasets, and support multiple types of statistical analysis (Cutler et al., 2007). RF methods extract static features described by a set of variables (*e*.*g*., the matrix of feature positions) present in the inputs, and process each feature independently without influences from temporally neighboring features (Lester et al., 2005; Nath et al., 2019). This intrinsic feature of RF classifiers can produce misclassifications that may not be easy to recognize (Sok et al., 2018). For example, a mouse quickly rearing during exploration may be indistinguishable from a long instance of standing or defending against an aggressor. This limitation of RF classifiers limits their utility for complex, temporally-evolving components of behavior.

Behavior intrinsically involves a sequential pattern of movements, and behavioral events occur in probabilistic relationships with the environment (Fountain et al., 2020). Hidden Markov models (HMMs), stochastic time-series models, infer that an observed event sequence is driven by a series of transitions between hidden states (Leos-Barajas et al., 2017). Once trained and validated, two types of information can be obtained from HMMs: the sequence of predicted hidden states and model transition probabilities between those states. HMMs have been extensively applied in multiple fields, such as speech recognition, bioinformatics, as well as the analysis of animal behavior (Findley et al., 2021; Jiang, 2021; Mor et al., 2021).

HMMs support behavioral classification tasks in many scenarios, but also have drawbacks. HMMs are unsupervised and parametric, but lack the human-interpretable labels associated with supervised ML methods (Bicego et al., 2006). Thus, one must explore and optimize many parameters to achieve results that have instructive value to human experimenters (Ahmadian Ramaki et al., 2018). For instance, the determination of the optimal number of hidden states is difficult to determine, often requiring subjective evaluation of results to avoid overgeneralization or fragmentation, both of which are barriers to interpretation and hypothesis testing (Deo, 2015; Wuest et al., 2016; Pohle et al., 2017; Liu and Shima, 2021). Despite their multiple advantages for quantifying temporally complex behaviors, the drawbacks of HMMs can limit their utility in behavioral neuroscience.

Given that both supervised and unsupervised ML algorithms have their own merits and demerits, several methods have employed both supervised and unsupervised ML methods, creating so-called hybrid ML models. Hybrid ML models attempt to use supervised and unsupervised methods (*e*.*g*., RFs and HMMs) to compensate for demerits of each process when used alone. In the present study, we used positional features obtained from DeepLabCut as inputs for a layered, RF-HMM hybrid ML analytic workflow (Fig. 1A). We employed this workflow to reveal that mice display fear-like response to 2,4,5-Trimethylthiazole (TMT) with a reduced tendency of approaching odor and a rapid-and-short investigative pattern. Also, we uncovered that predatory chemosensory cues resulted in an increased behavioral trend of risk assessment in mice. We found that the resulting layered, hybrid workflow produces rich, interpretable models that support hypothesis testing in chemosensory and other complex behavioral contexts.

**Figure 1.**
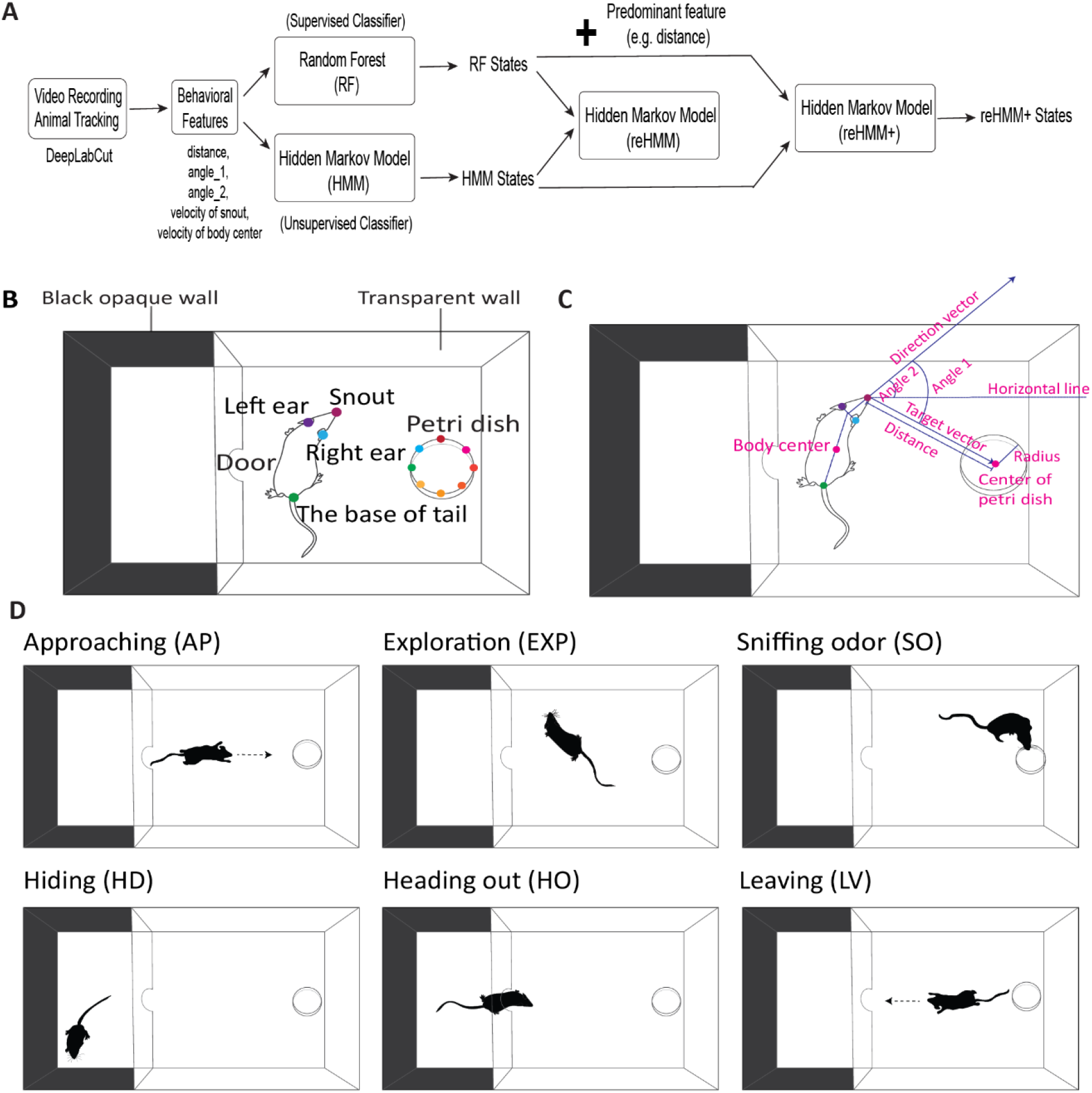
**A**, Architecture of analytic workflow and behavioral experiment overview. Mouse movement and body parts are tracked by using DeepLabCut software. Behavioral features (e.g., distance, angle_1, angle_2, and velocities of snout and body center) are calculated using DeepLabCut outputs and are used to train a Random Forest (RF) and a hidden Markov model (HMM) with equal numbers of states, separately. The per-frame predictions from RF and HMM are passed to a second HMM layer (reHMM). The predictions from RF and HMM plus predominate positional features are used to train a third HMM (“reHMM+”). **B**, Diagram of the behavioral test arena and DeepLabCut labeling. Mice are tracked by an overhead camera during video recording. For DeepLabCut labeling, four mouse body parts (snout, left ear, right ear, and the base of tail) are labeled. Eight labels equally spaced on a circle are used to label the petri dish. **C**, Graphical representation of derivative features. **D**. Ethogram including six behavioral states for mouse risk assessment behavior, including approaching (AP), exploration (EXP), sniffing odor (SO), hiding (HD), Heading out (HO), and Leaving (LV).

## Materials & Methods

### Mice

All animal procedures were in accordance with the University of Rochester University Committee on Animal Resources (UCAR) and the University of Texas Southwestern Medical Center Institutional Care and Use Committee (IACUC). Wild-type C57Bl/6 mice were obtained from the Jackson Laboratory (Stock #000651) or the Mouse Breeding Core of the University of Texas Southwestern Medical Center. All mice aged 8-15 weeks were kept with 12 h light/dark cycle (lights on from noon until midnight). Mice were given *ad libitum* access to food and water. Throughout the manuscript, the number of animals is described in text and figure legends.

### Behavioral test setup

Mouse risk assessment behavior was investigated in a customized two-chamber arena (Fig. 1B). The testing arena consisted of a small rectangular chamber with three opaque black walls [6” (L)×8” (W)×8” (H)] and a large rectangular chamber with three transparent walls [11” (L)×8” (W)× 8” (H)]. The two chambers were divided by a transparent wall with an open door. Mice could freely cross two chambers through the door. Mice are tracked by an overhead FLIR Blackfly camera (BFS-U3-16S2C-CS, USB 3.1) at 60 frames per second (FPS).

### DeepLabCut tracking

DeepLabCut (2.2.0.3) was used to track all points of interest (Mathis et al., 2018a; Nath et al., 2019), including four labels in the mouse body (snout, right ear, left ear, and the base of tail) and eight labels equally spaced in the petri dish (Fig. 1B). To create a robust network, we added a different number of new videos (frames) to retrain the existing network in each behavioral test because of slight discrepancies in experimental conditions, including lighting, testing arena, animal, and background. Cumulatively, a total of 9621 labeled frames were used to train ResNet-50-based neural networks with default parameters for 200,000-800,000 iterations. For all labeled frames, 80% was used for network training, whereas the remaining 20% was used for network evaluation. If the test error with p-cutoff was around 5 pixels (image size was 1160 by 716 pixels; 1 mm ≈ 2 pixels), this network was then used to analyze videos recorded in similar experimental conditions.

### Behavioral features and ethogram

The output of DeepLabCut was in the form of X/Y pixel coordinates of each label. We calculated several derivative features from DeepLabCut outputs that matched the experimental design (Fig. 1C). Specifically, we calculated the distance between the mouse snout and the center of the petri dish (“distance”). “Direction vector” was the vector from the midpoint of two ears towards the mouse snout. “Target vector” was the vector from the mouse snout to the center of the petri dish. “Angle_1” was the angle between the direction vector and the target vector. “Angle_2” was the angle between the direction vector and a horizontal line (X-axis). “Velocity of snout” was the instantaneous velocity (in pixels) of the snout from one frame to the next. “Body center” was the midpoint between the base of tail and the midpoint of two ears. “Velocity of body center” was the instantaneous velocity (in pixels) of the body center from one frame to the next.

To study mouse risk assessment behavior, we defined a simple ethogram consisting 6 behavioral states (Fig. 1D). Hiding (HD) was the state that the mouse stayed in the smaller, dark rectangular chamber. Heading out (HO) was the state that the mouse body crossed the door and towards the larger clear rectangular chamber. Approaching (AP) was the state that the mouse moved towards the petri dish directly from the small rectangular chamber. Leaving(LV) was the state that the mouse moved towards the small, dark rectangular chamber directly after sniffing the petri dish. Sniffing odor (SO) was the state that the mouse snout was located within the diameter of the petri dish.

### Odor stimuli preparation

2,4,5-Trimethylthiazole (TMT) was purchased from Sigma-Aldrich (St. Louis, MO, USA). TMT was diluted 1:9 (10%) with mineral oil (Saito et al., 2017). Female mouse urine was collected as previously described and stored at −80°C in a freezer (Holekamp et al., 2008) (“Female_urine”). Snake fecal samples were collected from the Department of Herpetology in the Dallas Zoo (project ID: DZ Project S2019-2). Fecal samples of inland taipan (*Oxyuranus microlepidotus*) and black mamba (*Dendroaspis polylepis*) were used in the behavior test. Snake feces was directly placed in the petri dish (“Feces”). For the preparation of snake fecal extracts, inland taipan feces particles (5 g) were placed in 50 mL of distilled water (dH_2_O). The feces mixture was homogenized by vortexing for 2 min and placed on ice on an orbital shaker overnight. On the second day, the feces mixture was homogenized by vortexing for 2 min and then sequentially subjected to two steps of centrifugations (10 mins at 2400xg at 4°C and 30 mins at 2800xg 4°C). The supernatant from two centrifugations was pooled in collection tubes (“s_unfiltered”) and filtered with a 0.22 μM filter (“s_filtered”). Fecal solids (“solid”) were also kept and stored at −80°C. Pure water (dH_2_O) was used as the control odor (“control”).

### Risk assessment behavior test

All behavioral tests were performed during the dark cycle under dim red light. For odor presentation, 3D-printed fake fecal particles (1 cm in length with an ellipse shape) were immersed in odor solutions overnight before the experiment day. Fake fecal particles (n = 5-7) were placed in the petri dish and kept wet during the experiment. To avoid cross-contamination, one petri dish was used only for one odor. In all behavioral tests, mice were exposed to each odor for 5 min.

For the first study, 24 C57Bl/6 male mice were randomly divided into 3 groups (n=8 for each group) and exposed to control, TMT, and female_urine, respectively. Without habituation, mice were placed in the test arena for 5 min. Behavioral tests were performed on three consecutive days and only one group (one odor) was tested each day.

For the second study, 16 C57Bl/6 male mice were used. Control, TMT, female_urine, and the feces were used as odor treatments. One day before the experimental test, mice were placed in the test arena for 20 min to habituate to the novel environment. On the testing day, mice were sequentially exposed to odors in sequence as “control-control-X-control-X-control-TMT”, in which X represented either female_urine or the feces in a random pattern. Because of its well-known fear-inducing effect, TMT was always the last odor delivered to mice. Interspersed control treatments were intended to reduce the potential for stimulus order effects. The second of the initial control treatments was used as the “control” sample for statistical purposes.

For the third study, 6 C57Bl/6 male mice were used. Control, inland taipan feces (Feces_1), black mamba feces (Feces_2), solid, s_unfiltered, and s_filtered stimuli were used as odor treatments. One day before the experimental test, mice were placed in the test arena for 20 min to habituate to the novel environment. Then, mice were subjected to two-day testing. On the first testing day, mice were sequentially exposed to odors in sequence as “control-control-X-control-X-control-X-control-Feces_1”, in which X was one of solid, s_unfiltered, and s_filtered in a random pattern. The testing on the second day exactly repeated the stimulus order of the first day, except for an additional “control-Feces_2” that was added at the very end. As above, interspersed control treatments were interspersed between odor treatments, and the second of the initial control treatments was used as the “control” for statistical analysis.

### Random forest

RF classification was performed by using Python (3.8.12) in the Conda environment (4.10.3). The RF classifier (sklearn.ensemble.RandomForestClassifier) was loaded from the Scikit-learn library (Pedregosa et al., 2011). The hyperparameter n_estimators was defined as 50 and all others were default values. As the ground truth (GT) in this study, 6 behavioral states (HD, AP, HO, LV, EXP, and SO) were manually annotated frame-by-frame in 12 videos. Five behavioral features (distance, angle_1, angle_2, velocity of snout, and velocity of body center) extracted from 12 manual annotated videos (a total of 149,258 frames) were used to train the RF model. For all frames, 80% was used for the RF classifier training, whereas the remaining 20% was used for model evaluation. The RF performance was evaluated by 10-fold cross-validation (sklearn.model_selection.KFold). Feature importance analysis was conducted by using the module (sklearn.feature_selection. SelectFromModel) from the Scikit-learn library. To further improve the performance of the RF, behavioral feature data were smoothed using a rolling median filter (window sizes= 5, 10, 15, 30, and 60) from the Pandas library (pandas.Series.rolling). After data smoothing, 30 behavioral features (5 original features plus 25 smoothed features) were used to train a new RF model. The same approaches mentioned above were used to evaluate the performance of the new RF model.

### Hidden Markov Models

Hidden Markov Models (HMMs) were implemented by using the model (hmm.GaussianHMM) from the hmmlearn Python library (https://github.com/hmmlearn/hmmlearn). The hyperparameters of the HMM model were as follows: covariance_type=“diag”, n_iter=100, verbose=True, random_state=0. “n_components” representing the number of hidden states was adjusted as needed. The same 5 behavioral features used to train the RF were also fed to train the HMM classifier. The outcomes of the HMM were evaluated by comparing it with the GT and manually checking the plots of the behavioral features of each HMM hidden state. For HMM model optimization, Expectation-Maximization (EM) algorithm and Bayesian Information criterion (BIC) were used as criteria to determine the “n_components” parameter. For the second HMM (“reHMM”) and third HMM (“reHMM+”), the training data sequences (RF classification, HMM classification, and distance) were compressed by replacing with the most frequent element (RF classification and HMM classification)/median (distance) in every 15 frames. “n_components” of reHMM and reHMM+ models were 6 and other hyperparameters were identical with HMM. The classifications of RF and HMM were fed to train the reHMM. The classifications of RF and HMM and distance were used to train the reHMM+. The same evaluation methods for HMM were used to evaluate the performance of the reHMM and reHMM+. The code and example data are available in a GitHub repository (https://github.com/julianmeeks/Wang_ThreatAssessment).

### Statistical analysis

Data analysis was performed using JMP Pro 16 (SAS Institute, Cary, NC, USA). Data are expressed as means ± standard error. Behavioral responses between different odor conditions was compared using a one-way ANOVA or Student’s *t*-test. *p* < 0.05 was regarded as a statistically significant difference.

## Results

### Optimizing and evaluating the RF classifier

We manually annotated 12 top-down videos of mouse behavior (149,292 frames) in our behavioral arena to serve as a “ground truth” (GT) dataset. Tracking data from DeepLabCut were distilled to five derivative features, including the distance from the snout to the petri dish, head angle relative to the sample petri dish center, head angle relative to the horizontal image axis, and the snout and body center velocities (see Materials and Methods, Fig. 1C). Behavioral features of the GT were plotted for frames annotated as belonging to 6 behavioral states of a custom ethogram for the experimental setup (AP, HO, EXP, HD, SO, and LV, see Materials and Methods, Fig. 2A).

**Figure 2.**
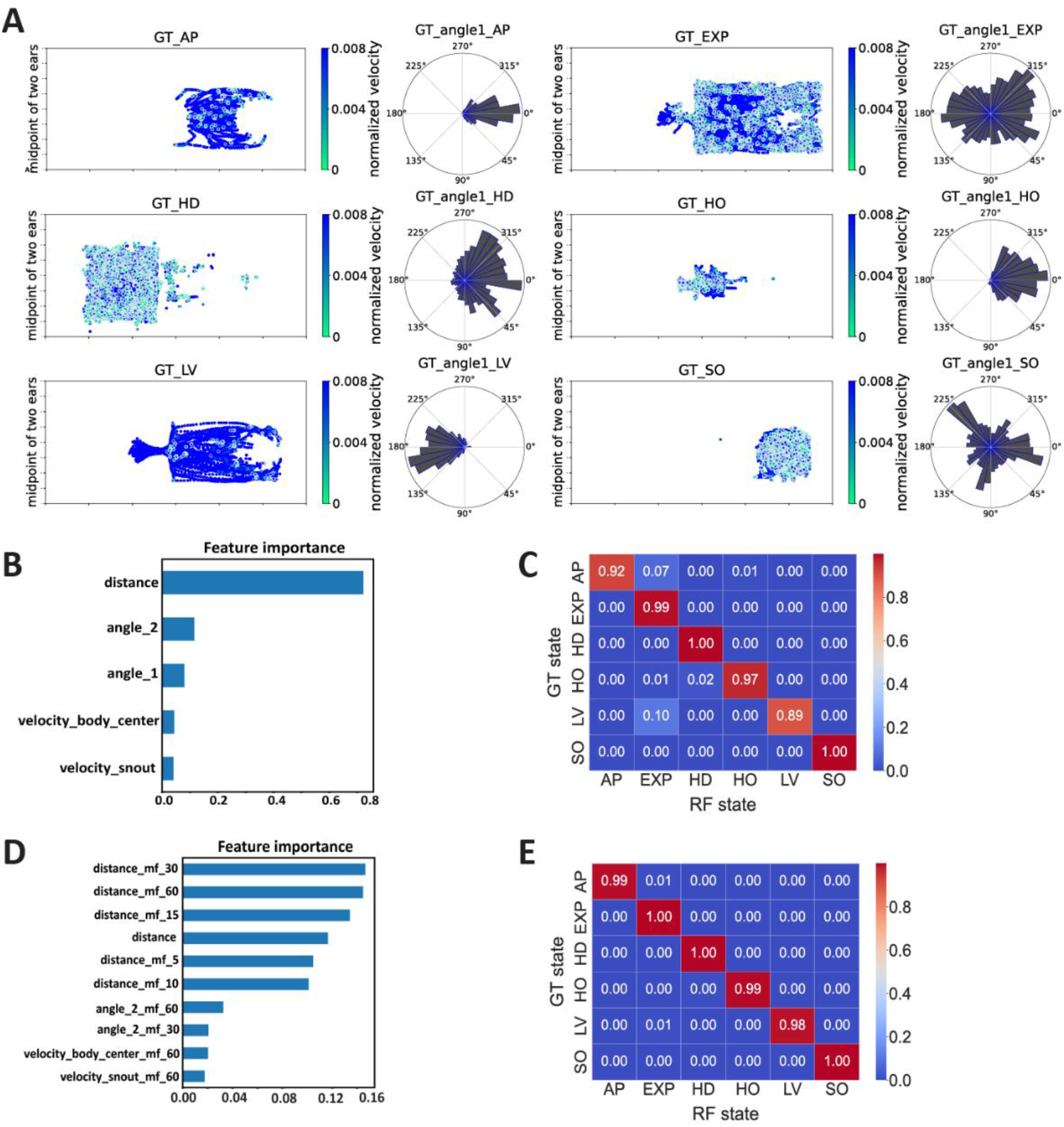
Plots of ground truth (GT) and performance of Random Forest (RF). **A**, Graphical representations of the GT behavioral states. Each dot denotes the mouse position (midpoint between the ears) during manually annotated frames, with color indicating the normalized instantaneous velocity of the animal center. At right are polar plots indicating the direction of the mouse head relative to the horizontal axis. **B**, Feature importance for the RF. **C**, Confusion matrix for the RF versus the GT. **D**, Feature importance of the optimized RF, which included rolling median filters of each derivative feature with temporal sliding windows (5, 10, 15, 30, and 60 frames) **E**, Confusion matrix for the optimized RF versus GT.

RF models have been widely applied in the classification of animal behavior due to their high classification accuracy and versatility (Valletta et al., 2017; Wang, 2019). We first evaluated the performance of the RF against GT data. We used the five derived features described above to train a RF classifier, achieving an overall accuracy of 0.9513. Feature importance analysis revealed that the distance between the snout and the sample petri dish was the most informative feature for the RF model (Fig. 2B). Despite the high overall accuracy of the RF, the accuracy for each behavioral state was variable, ranging from 0.89 to 1.00 (Fig. 2C). Approximately 10% of “leaving” (LV) and 7% of “approach” (AP) were misclassified as “exploration” (EXP). These misclassifications may have been a result of fewer training images for these states compared to others, as both LV and AP were transient states occupying many fewer overall video frames compared to “hiding” (HD) and EXP. Another potential cause might be the inherent limitations of RF models, specifically their blindness to temporal features of the dataset. RF models distinguish class boundaries independently of neighboring data (*i*.*e*., timepoints before and after the analyzed frame), leading to a lack of temporal relations between outcomes (Rubinstein and Hastie, 1997; Lester et al., 2005). In an attempt to provide the model with some temporal information, we temporally smoothed the data with rolling median filters of variable window sizes (James et al., 2016; Cook et al., 2019) (Fig. 2D). After inclusion of temporally filtered features, distances between the snout and petri dish remained the most informative of the RF classification (Fig. 2D). This also improved the overall accuracy to 0.9931. Remarkably, the classification accuracy for AP and LV reached 0.99 and 0.98, respectively (Fig. 2E). These results show that RF models achieve high classification accuracy in these conditions, and that RF models benefit from the inclusion of temporally smoothed data.

### Optimizing and evaluating the HMM classifier

Though the RF model achieved high classification accuracy, its inherent limitations related to temporal components of behavior were of some concern (Sok et al., 2018). Hidden Markov Models (HMMs) are another popular ML method for assessing highly dynamic behavioral data (Kim et al., 2009; Liu et al., 2015). HMMs infer transitions between hidden (latent) states that best predict observed data, potentially compensating for drawbacks of RF models. As an initial test of utility, we compared the performance of RFs and HMMs directly (Fig. 3). The same five derivative features used for the RF models shown in Fig. 2 were used to train HMMs (Fig. 3B-D). When given an equal number of latent states to the RF (6), the HMM identified some states that roughly matched GT states (especially “sniffing odor” [SO] and “leaving” [LV], Fig. 3B).However, most HMM states were not clearly interpretable (Fig. 3A), and the confusion matrix for the 6-state HMM showed that this approach greatly differed from the supervised, more human-interpretable RF (Fig. 3B). Because the number of latent states is generally defined by users, we generated HMMs with increasing numbers of hidden states in two batches, ranging from 6-12 states (Fig. 3-1A-B) and 13-24 states (Fig. 3-2A-B). The HMM models with the best performance in these tests (HMMs with 11 and 23 states, respectively) had confusion matrixes that were no more interpretable than the 6-state model (Fig. 3C-D, Fig. 3-1C, Fig. 3-2C). Thus, despite advantages related to sequential information, HMMs missed considerable, human-interpretable information within these datasets.

**Figure 3.**
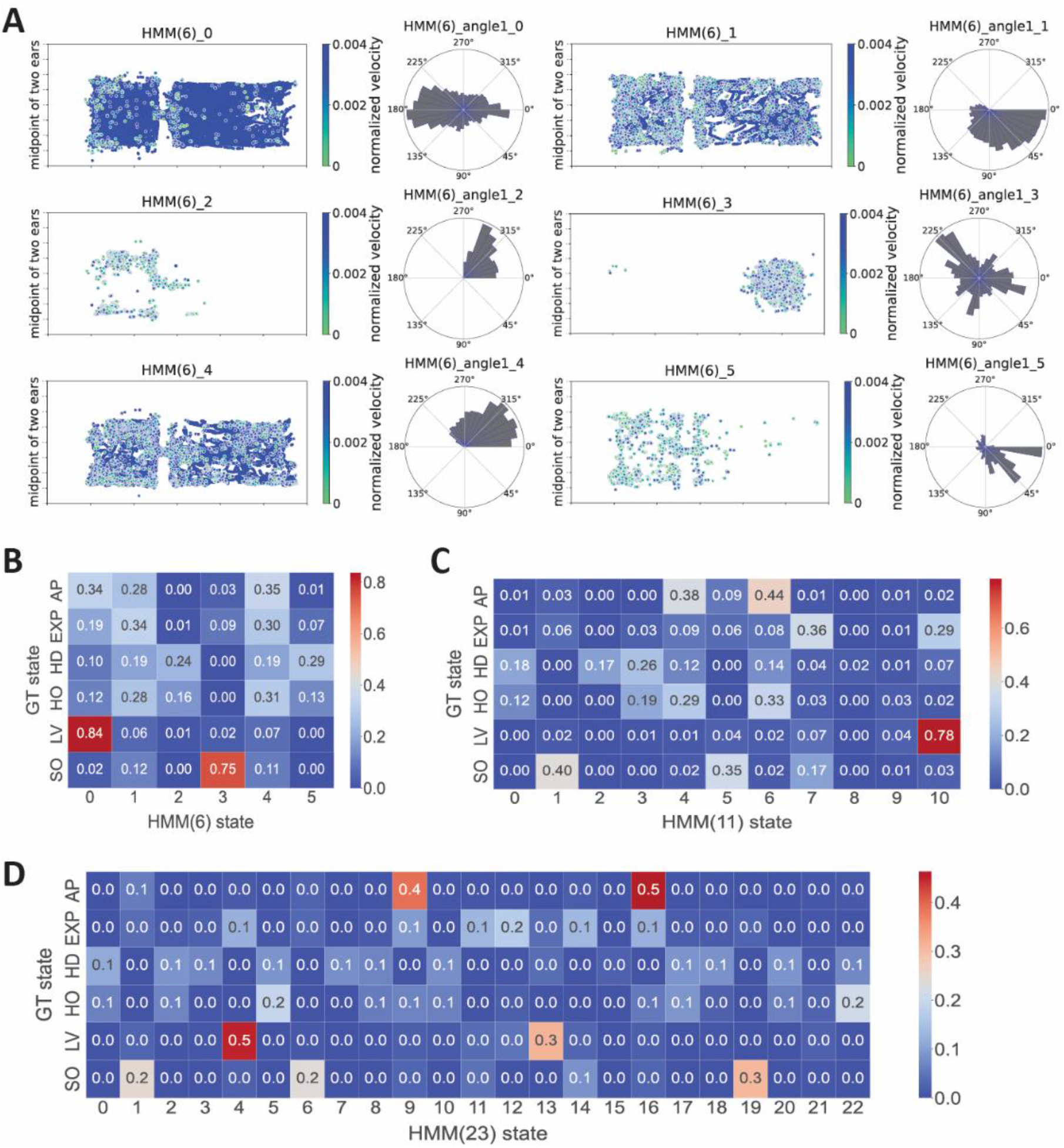
Performance of the hidden Markov model (HMM). **A**, Graphical representations of HMM behavioral states for a 6-state classification (HMM(6)). Each dot denotes the mouse position (midpoint between the ears) during manually annotated frames, with color indicating the normalized instantaneous velocity of the animal center. At right are polar plots indicating the direction of the mouse head relative to the horizontal axis. **B**, Confusion matrix for the HMM(6) versus the GT. **C**, Confusion matrix for the HMM for 11-state classification (HMM(11)) versus the GT. **D**, Confusion matrix for the HMM for 23-state classification (HMM(23))versus the GT.

**Figure 3-1.**
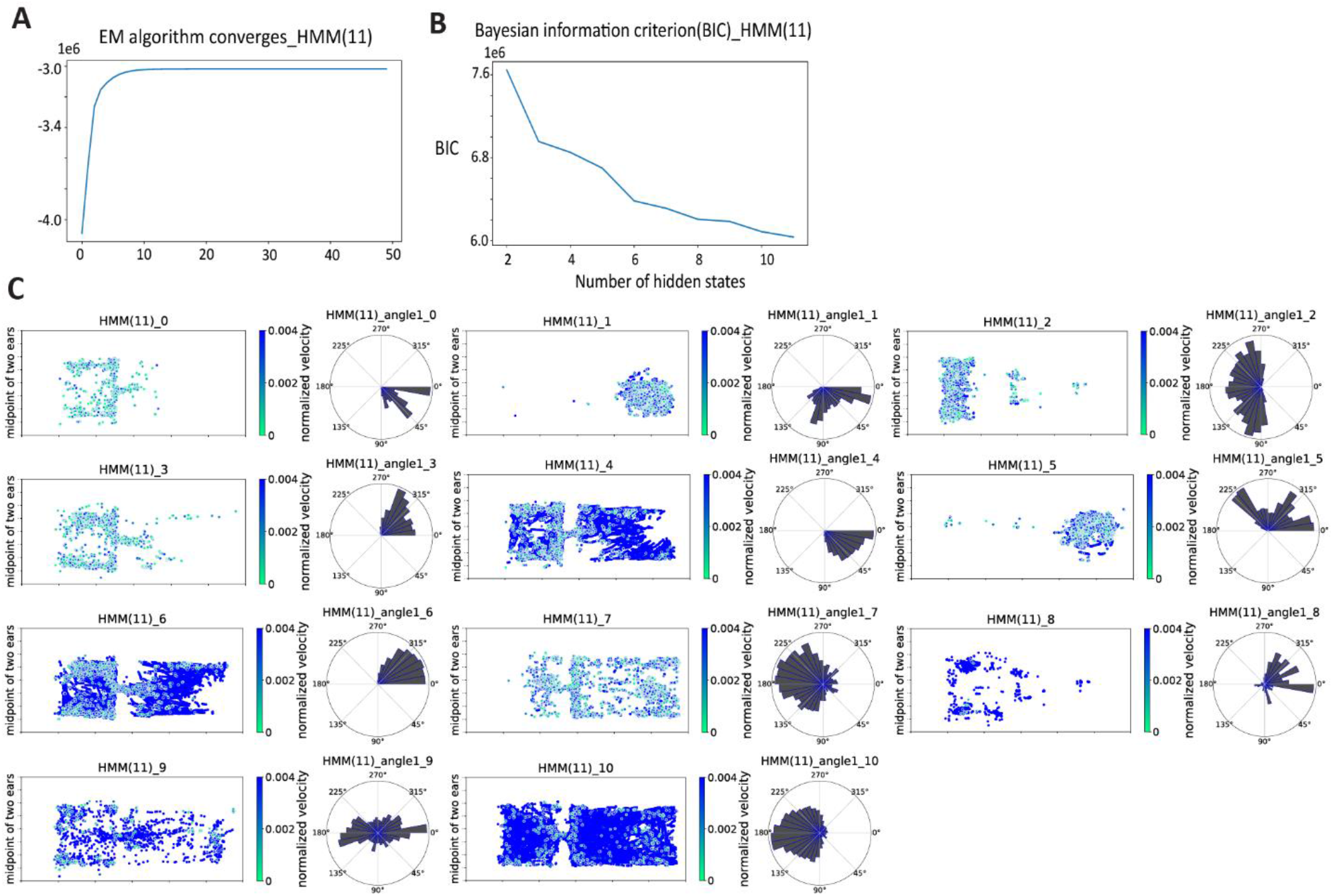
Overview of hidden Markov model for 11-state classification. **A**, Expectation-Maximization (EM) algorithm for hidden Markov model for 11-state classification (HMM(11)) relative to the number of training iterations. **B**, Bayesian Information Criterion (BIC) scores of HMM(11) relative to the number of hidden states. **C**, Graphical representations of hidden behavioral states, as predicted by HMM(11). For each behavioral state, dots represent the midpoint of two ears, and the color represents the velocity of the body center. The right polar plot represents the angle between the head direction vector and the horizontal x-axis.

**Figure 3-2.**
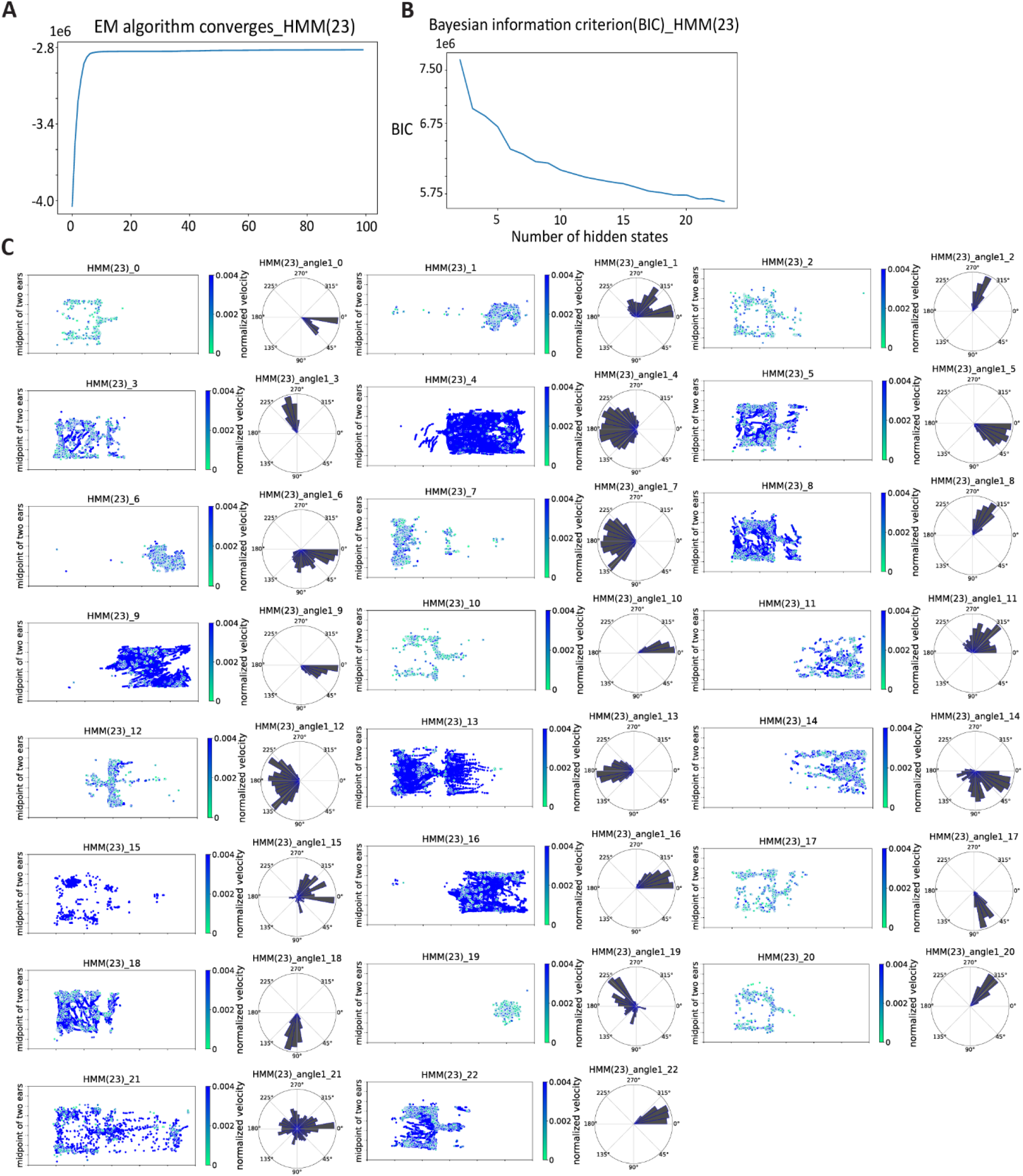
Overview of hidden Markov model for 23-state classification. **A**, Expectation-Maximization (EM) algorithm for hidden Markov model for 23-state classification (HMM(23)) relative to the number of training iterations. **B**, Bayesian Information Criterion (BIC) scores of HMM(23) relative to the number of hidden states. **C**, Graphical representations of hidden behavioral states, as predicted by HMM(23). The dots represent the midpoint of two ears, and the color represents the velocity of the body center. The right polar plot represents the angle between the head direction vector and the horizontal x-axis.

### Hybrid modes reHMM and reHMM+

Given the limitations of RF and HMM strategies, we next considered a hybrid ML models, which have been demonstrated to outperform either alone (Lester et al., 2005; Antos et al., 2014; Sok et al., 2018). We first used the outputs of RF and HMM to train a new HMM classifier for 6-state classification, named “reHMM” (Fig. 4). reHMM model output showed a closer match between reHMM states and GT compared to HMM-alone (Fig. 4A-B). For example, the reHMM states 0 and 2 together split frames associated with the GT HD state, accounting for 52% and 48%, respectively (Fig. 4B). The reHMM state 5 included 64% of AP and 56% of HO, which share the feature that the mouse body is oriented towards odorant cues.

**Figure 4.**
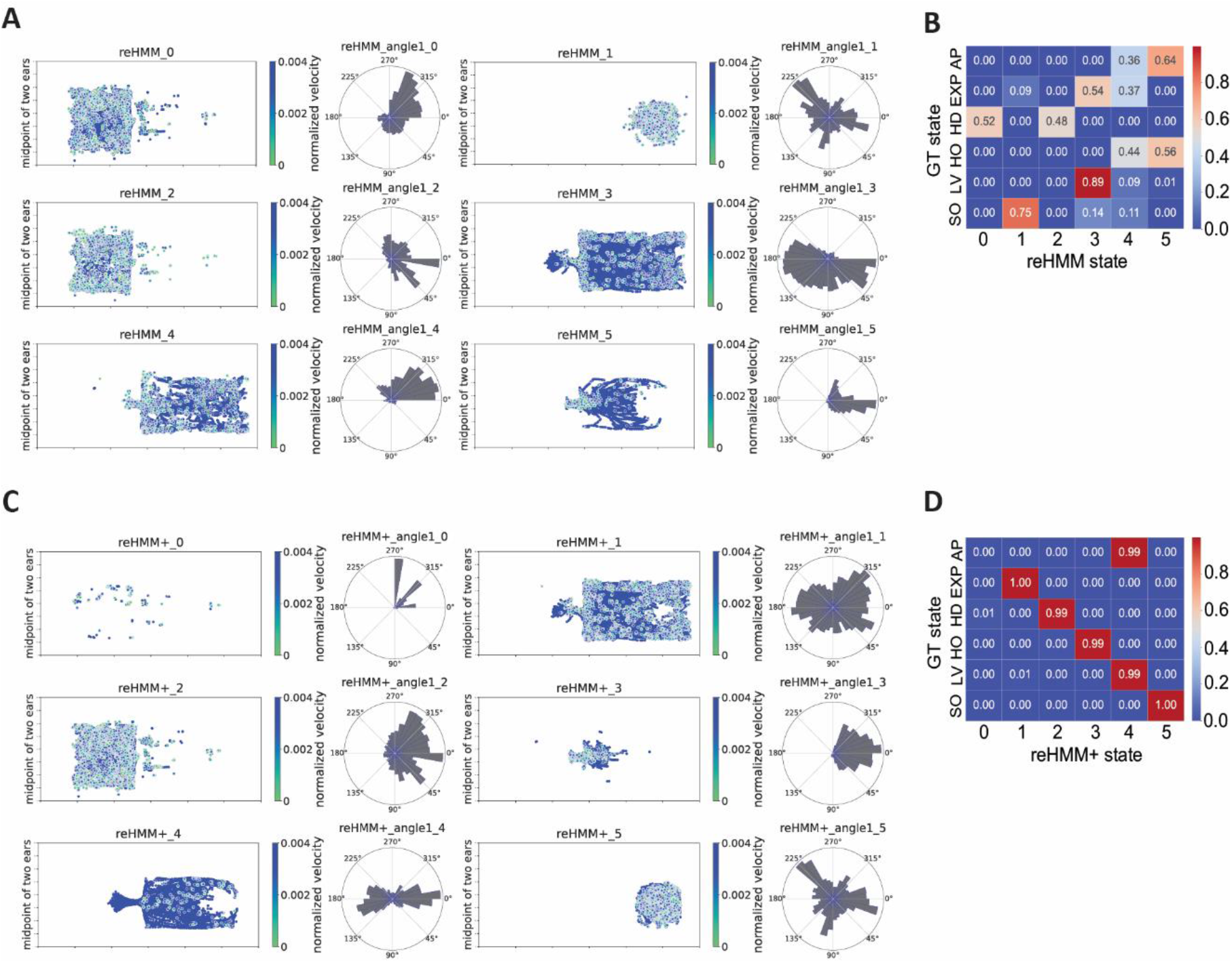
Performance of reHMM and reHMM+. **A**, Graphical representations of hidden behavioral states, as predicted by the reHMM. Dots represent the midpoint of the two ears and the color represents the velocity of the body center. Polar plots represent the angle between the head direction vector and the horizontal x-axis. **B**, Confusion matrix for reHMM versus the GT. **C**, Graphical representations of hidden behavioral states, as predicted by the reHMM+. **D**, Confusion matrix for reHMM+ versus the GT.

Given the improvement of reHMM compared to the 6-state HMM alone, we sought to further enhance the reHMM model’s interpretability. Leontjeva and Kuzovkin reported that combining dynamic and static features enables classification models to simultaneously capture static information and temporal dynamics (Leontjeva and Kuzovkin, 2016). In our effort to optimize the RF, the distance between the mouse snout and the stimulus order was the most informative static feature, so we added this feature to the RF-only and HMM-only predictions (creating a three-dimensional matrix). This input matrix, including the predictions of RF-only, HMM-only, and the distance measurement, was used to train a reHMM variant, named “reHMM+” (Fig. 4C-D). reHMM+ states were highly interpretable and mapped strongly to GT states, suggesting similar classifying strength of the reHMM+ to the RF model (Fig. 4C-D). For example, the reHMM+ state 4 contained both AP and LV, which could be regarded as a back-and-forth state. SO state was exclusively distributed in the reHMM+ state 5, providing an avenue to decipher SO patterns and transition frequencies between SO and others. Overall, this layered, hybrid reHMM+ model showed the capacity to classify behaviors in these experimental conditions with high interpretability and accuracy, incorporating static and temporal features to achieve its predictions.

### Behavioral outputs of male mice in response to TMT and female mouse urine

We next sought to test the reHMM+ model to evaluate its performance in mouse risk assessment behavioral assays (Fig. 5). To simplify interpretation, the reHMM+ states representing HO, AP, and LV were merged into one state reHMM+ state 2. This state included transition states between the entry to the safe area and the odor object, essentially a back-and-forth (B-A-F) state. State 0, 1, and 3 effectively encapsulated HD, EXP, and SO, respectively. For visualization, each reorganized reHMM+ state was also assigned a color code; red represented avoidant HD and yellow represented neutral EXP, respectively. Orange represented B-A-F state and green represented attractive SO, respectively (Fig. 5A).

**Figure 5.**
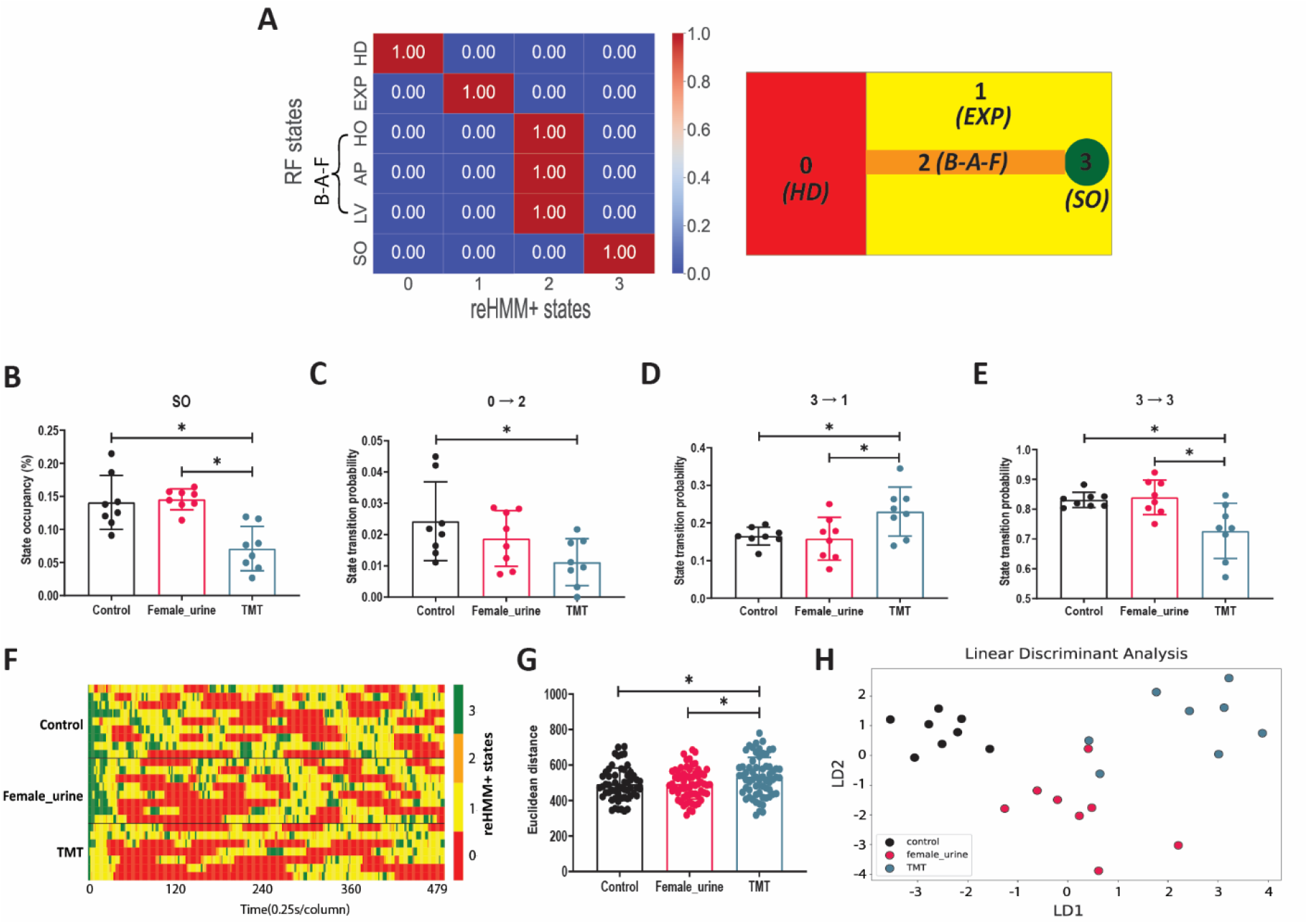
Behavioral responses of male mice to TMT and female mouse urine. Analysis was conducted for a 2 min window starting with the first SO event. A, Confusion matrix for the merged reHMM+ states versus the RF states. For simplicity, the three reHMM+ states matching the RF states approaching (AP), heading out (HO), and Leaving (LV) were merged into the back-and-forth (B-A-F) state. Right: graphical representation of the interpretation of merged reHMM+ states. Red indicates state 0 (hiding(HD)). Yellow indicates state 1 (exploration(EXP)). Orange indicates state 2 (back-and-forth(B-A-F)). Green indicates state 3 (sniffing odor (SO)). B, Occupancy analysis for the reHMM+ state SO. C-E, Transition probabilities between listed reHMM+ states F, Heatmap illustrating behavioral sequence aligned to the first SO event. Each row indicates one mouse. Each column indicates the time (0.25s). The color code is identical to the description in panel A. G, Behavioral sequence similarity, as evaluated by Euclidean distance. H, Plot of linear discriminant analysis. * indicates p < 0.05. For this experiment, 24 mice (n=8 for each treatment group) were available for analysis.

In an initial proof-of-concept experiment, TMT, female mouse urine, and pure water were used as test odorants. TMT, a compound derived from red fox feces, has been widely used to induce fear-like behavioral responses in rodents, such as freezing, avoidance, and defensive burying (Fendt et al., 2005). Consistent with previous studies, the reHMM+ output showed that mice spend less time on investigating TMT compared to female urine and pure water (Fig. 5B). Moreover, reHMM+ revealed that mice had a lower probability of transiting from state 0 to 2 and state 3 to 3, while a higher probability of switching from state 3 to 1 in the presence of TMT (Fig. 5C-E). These observations suggest that TMT-exposed mice were inclined to stay in HD and leave SO. During the first 2 min following the first SO event (the first close investigation of the test odorant), the average intervals between two consecutive SOs were larger in TMT-treated mice compared with pure water (Fig. 5-1A). Also, the average duration of SO was much shorter for mice encountering TMT compared with animals faced with water and female urine (Fig. 5-1B). These data indicate that mice displayed less frequent and shorter investigative responses to TMT. On the other hand, we did not find differences in the total number of SO events, the latency to the first SO event, or total movement distance between the three treatments (Fig. 5-1C-E). We next measured the Euclidean distance between the first-SO-aligned reHMM+ classifications to investigate the behavioral sequence similarity in each odor treatment, finding that the behavioral sequence of TMT-treated mice differed from those of mice treated with pure water and female urine (Fig. 5F-G). Because so many metrics were produced by this analytical workflow, including reHMM+ state occupancy, the transition probability of reHMM+ states, SO frequency, SO duration, SO latency, the total number of SO, total movement distance, and behavioral sequence similarity. In order to compare the overall behavioral difference across all analytic metrics, we utilized linear discriminant analysis (LDA), which identifies multidimensional axes (Eigenvectors) that best separate experimental groups. LDA analysis revealed that the behavioral outputs from mice confronted with different odor cues could be readily distinguished (Fig. 5H). These data show that the reHMM+ analytic workflow is well-suited for behavioral pattern analysis in response to appetitive and aversive odorants.

**Figure 5-1.**
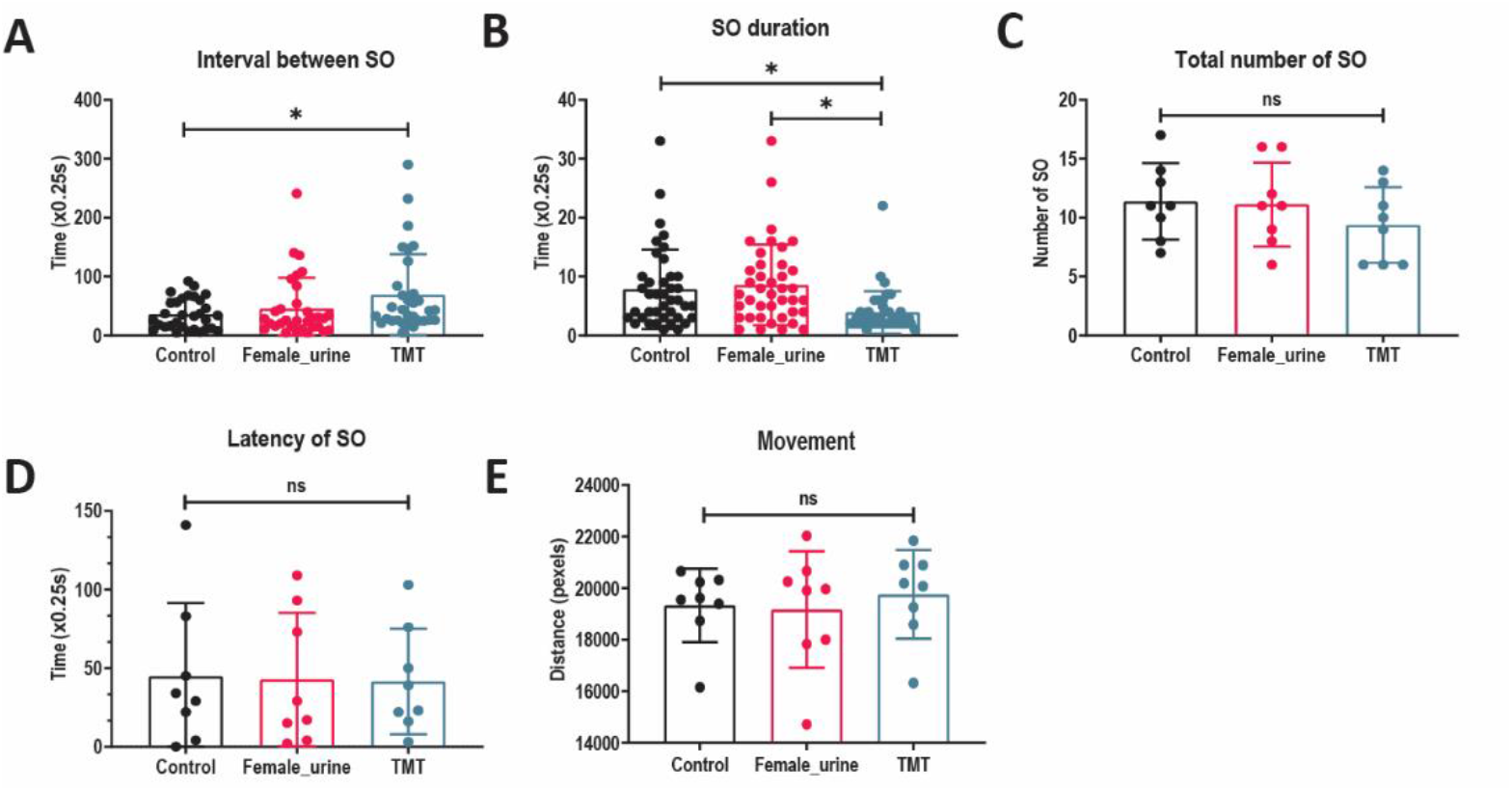
Additional analysis of male mouse behavioral responses to TMT and female mouse urine. The **A**, Interval between two consecutive SOs. **B**, Duration of each SO. **C**, Total number of SO. **D**, Latency of the first SO. **E**, Total movement distance. * indicates *p* < 0.05. For this experiment, 24 mice (n=8 for each treatment group) were available for analysis.

### Behavioral outputs of male mice in response to TMT, female mouse urine, and snake feces

Quantifying threat assessment behavior can be a challenging task, especially in the context of complex chemosignal blends that laboratory strains of mice have never encountered in their natural context (e.g. predator cues). We investigated the mouse behavioral responses to snake feces, a predatory odor cue. In this experiment, TMT, female mouse urine, and pure water were used as negative, positive, and neutral controls, respectively. In this experiment, each animal was exposed to all 4 odorants in a pseudorandom order (the exception was that negative control was always first and TMT always last).

reHMM+ state occupancy analysis revealed that snake feces-treated mice spent more time in SO state than TMT, whereas less time compared with female urine (Fig. 6A). Similar to the observations of the first pilot study, exposure to TMT was associated with a higher occupancy of B-A-F, but a lower level of occupancy in SO, compared with female urine (Fig. 6A-C). Mice investigating female mouse urine displayed higher occupancy of SO while lower HD relative to pure water (Fig. 6A-B). reHMM+ demonstrated that snake feces caused a higher probability of transiting from state 1 to 3 compared with pure water (Fig. 6D). Compared with TMT, snake feces-encountered mice displayed a lower probability of leaving state 3 (3 to 1) (Fig. 6E). Interestingly, the probability of staying in state 3 (aligned with the “sniffing odor” RF condition) in response to snake feces was higher than TMT, but lower than female urine (Fig. 6F).

**Figure 6.**
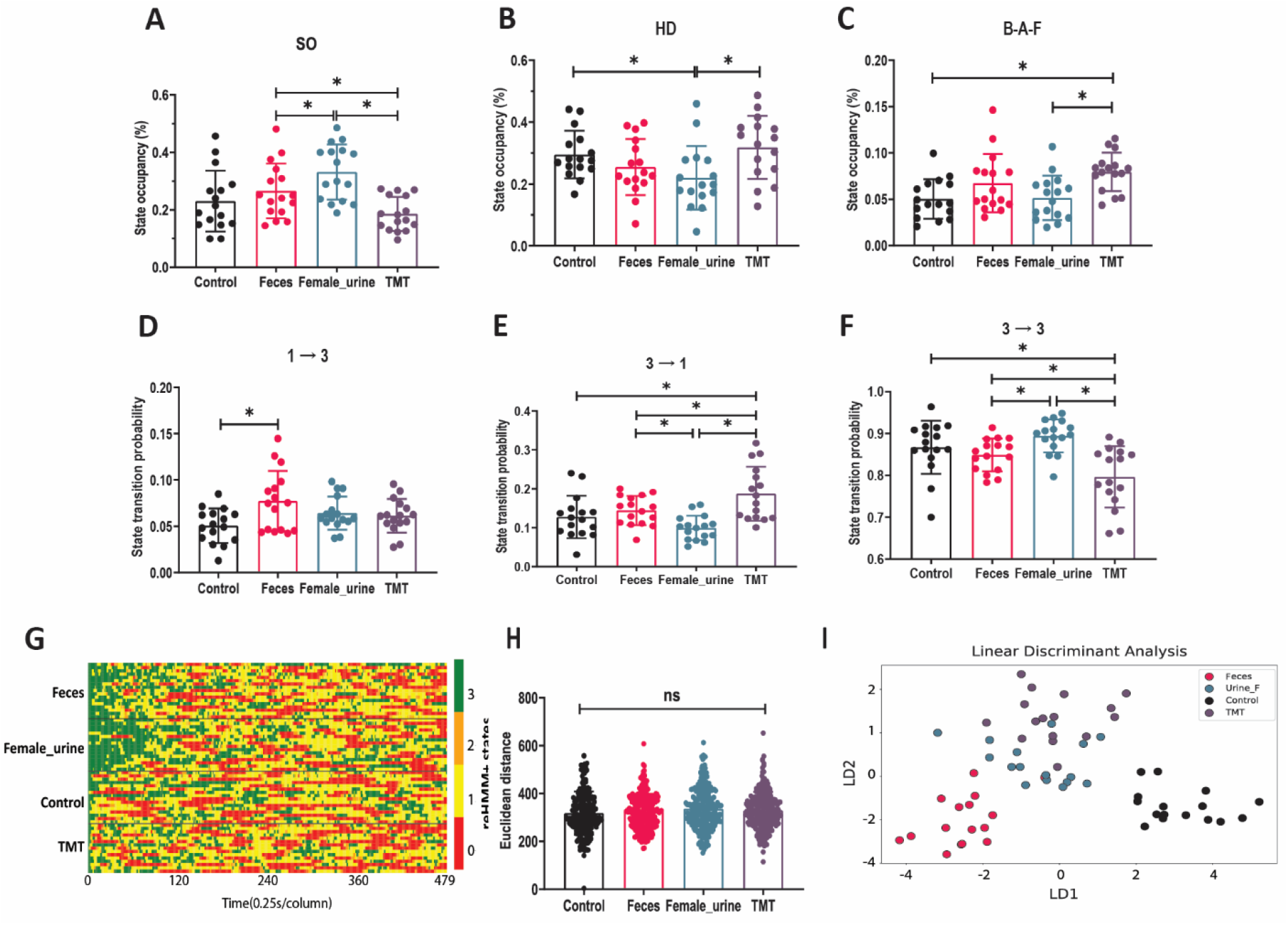
Male mouse behavioral responses to female mouse urine, snake feces, and TMT. **A-C**, Occupancy analysis for listed reHMM+ states. **D-F**, Transition probabilities between listed reHMM+ states. **G**, Heatmap illustrating behavioral sequences aligned to the first SO event. Each row indicates one mouse. Each column indicates the time (0.25s). Red indicates state 0 (hiding(HD)). Yellow indicates state 1 (exploration(EXP)). Orange indicates state 2 (back-and-forth(B-A-F)). Green indicates state 3 (sniffing odor (SO)). **I**, Behavioral sequence similarity, as evaluated by Euclidean distance. **J**, Plot of linear discriminant analysis. * indicates *p* < 0.05. For this experiment, 16 mice were available for analysis.

Also, this analysis suggested that snake feces caused a higher frequency of SO (shorter SO interval) and more total number of SO relative to pure water (Fig. 6-1A, C). This suggested that the animals were actively assessing the snake feces, and that they found it mildly threatening compared to conspecific cues. In these conditions, TMT treatment was associated with increased overall SO frequency, but an overall decreased SO duration compared with pure water (Fig. 6-1A-D). This suggested that in these conditions, where mice are habituated to their environment by a series of control and odorant presentations, TMT is not as overtly fear-inducing as it is in other conditions. As expected, female mouse urine caused the longest SO duration among all treatments (Fig. 6-1B). In these conditions, we observed no difference in the total movement, suggesting that animals did not freeze for long periods of time in these conditions, even in the presence of the well-established aversive TMT odorant (Fig. 6-1E).

**Figure 6-1.**
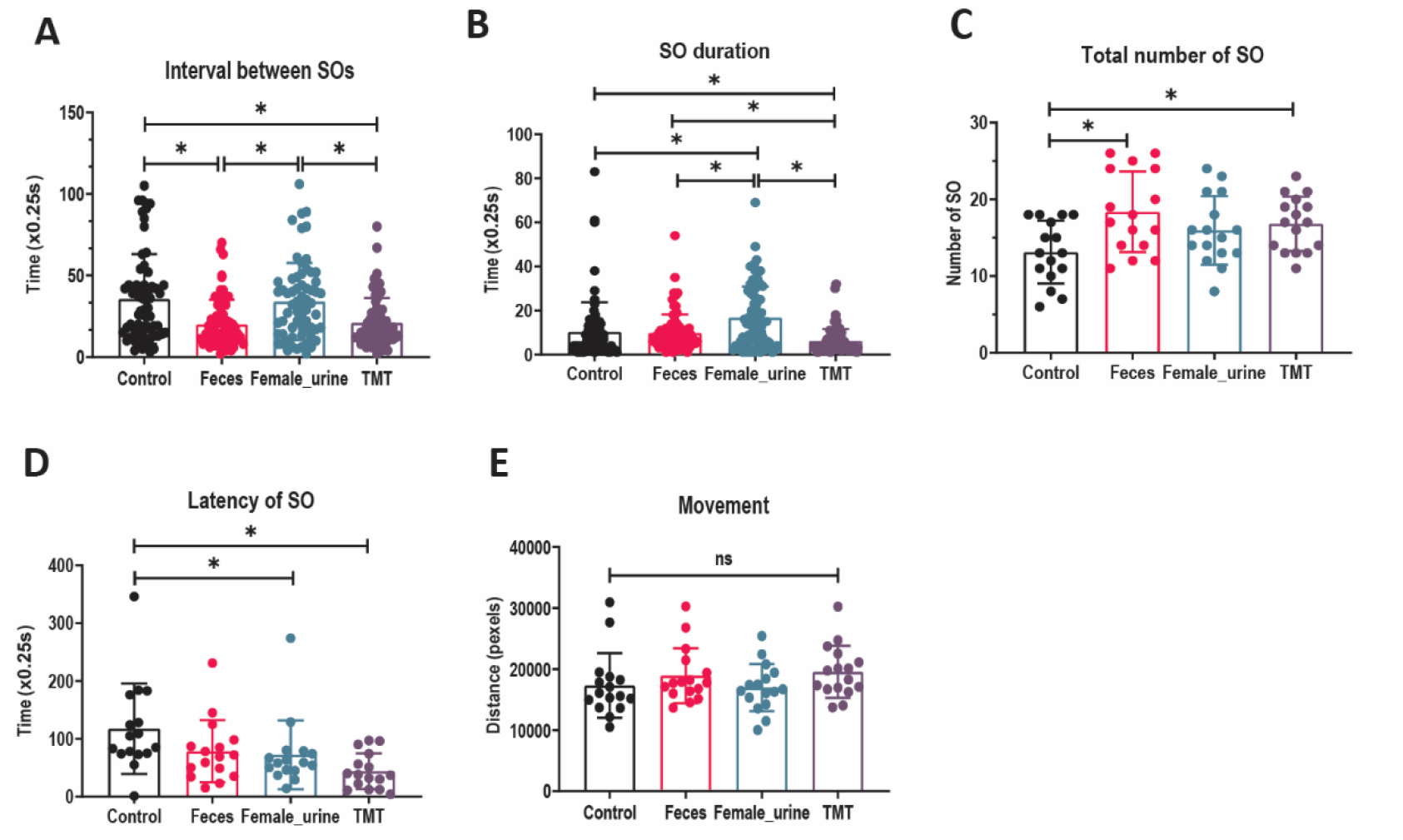
Additional analysis of male mice behavioral responses to female mouse urine, snake feces, and TMT. **A**, Interval between two consecutive SOs. **B**, Duration of each SO. **C**, Total number of SO. **D**, Latency of the first SO. **E**, Total movement distance. * indicates *p* < 0.05. For this experiment, 16 mice were available for analysis.

First-SO-aligned behavioral sequence analysis indicated that the behavioral responses to all test and control odorants were not generally distinguishable from each other (Fig. 6H-I). LDA analysis, on the other hand, which incorporated a broader range of metrics, revealed that behavioral responses to snake feces were separable from other odorants, including TMT (Fig. 6J). These studies show that mice exposed to multiple odorant presentations in this threat assessment assay respond differently to odorants than when they encounter them naively. The data also suggest that mice respond to novel predatory cues with an increased behavioral trend of risk assessment, not overt aversion.

### Behavioral outputs of male mice in response to snake feces and its extracts

A major goal in chemosensory neuroscience is to identify specific chemosignals that drive behavioral changes. To support the eventual identification of novel predatory chemosignals, we introduced animals to fractions of snake feces, including unfiltered and sterile filtered extracts, as well as residual solids (Fig. 7). Snake feces and pure water were used as positive and negative controls, respectively. A second snake fecal treatment from a separate species was used in place of TMT as the final positive control stimulus. Model outputs showed that filtered and unfiltered extracts and the remaining solid caused higher SO occupancy than pure water, but less time spent in SO compared to snake feces (Fig. 7A). Both snake feces treatments were associated with a lower HD occupancy and probability of staying in state 0, compared with pure water (Fig. 7B-C). Filtered and unfiltered extracts induced a higher probability of transiting from state 1 to 3 and from state 0 to 2, compared with pure water (Fig. 7D-E), whereas mice treated with filtered and unfiltered extracts displayed a higher probability of staying in state 1 relative to two snake feces (Fig. 7F). Similar to the study shown in Figure 6, snake feces increased the total number of SO events compared with pure water (Fig. 7-1B-C). Unfiltered extracts also increased SO events and SO duration, while filtered extracts only increased the total number of SO (Fig. 7-1B-C). No difference in SO interval, the latency of SO and movement distance was observed (Fig. 7-1A, D, E). The first-SO-aligned behavioral sequences of filtered and unfiltered extracts and the remaining solid differed from pure water and two snake feces (Fig. 7G-H). Finally, LDA revealed that the behavioral outputs of all odorants could be distinguished, with the exception of the remaining solid and pure water control conditions (Fig. 7I). In all, these results suggest filtered and unfiltered extracts induced quantitatively different risk assessment behaviors than feces, suggesting that behavioral responses to complex odorant mixtures depend on the specific chemosignals present, not the presence or absence of a single component of the mixture.

**Figure 7.**
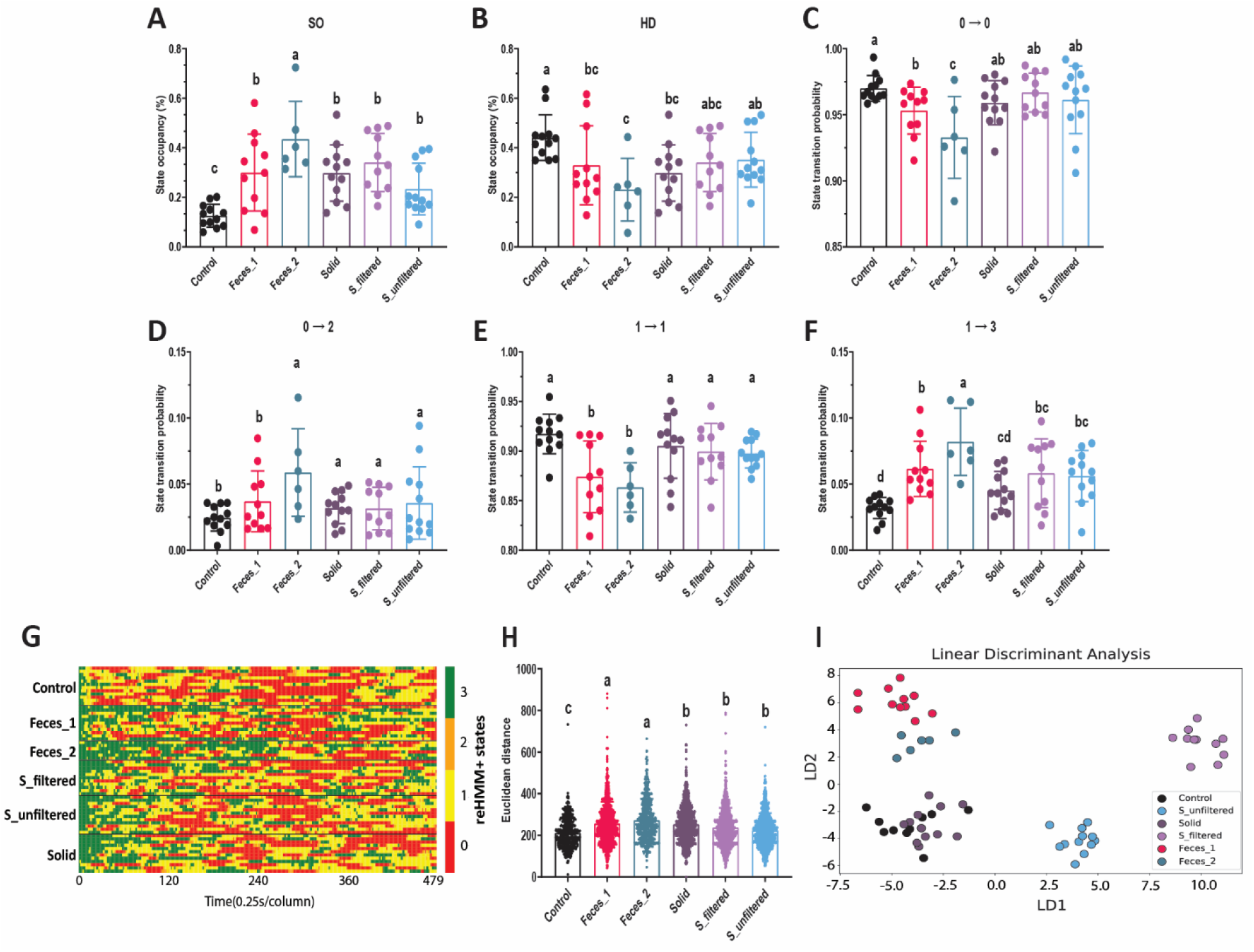
Male mouse behavioral responses to snake feces and its extracts. **A-B**, Occupancy analysis for listed reHMM+ states. **C-F**, Transition probabilities between listed reHMM+ states. **G**, Heatmap illustrating behavioral sequence. Each row indicates one mouse. Each column indicates the time (0.25s). Red indicates state 0 (hiding(HD)). Yellow indicates state 1 (exploration(EXP)). Orange indicates state 2 (back-and-forth(B-A-F)). Green indicates state 3 (sniffing odor (SO)). **H**, Behavioral sequence similarity, as evaluated by Euclidean distance. **I**, Plot of linear discriminant analysis. * indicates *p* < 0.05. For this experiment, 6 mice were available for analysis.

**Figure 7-1.**
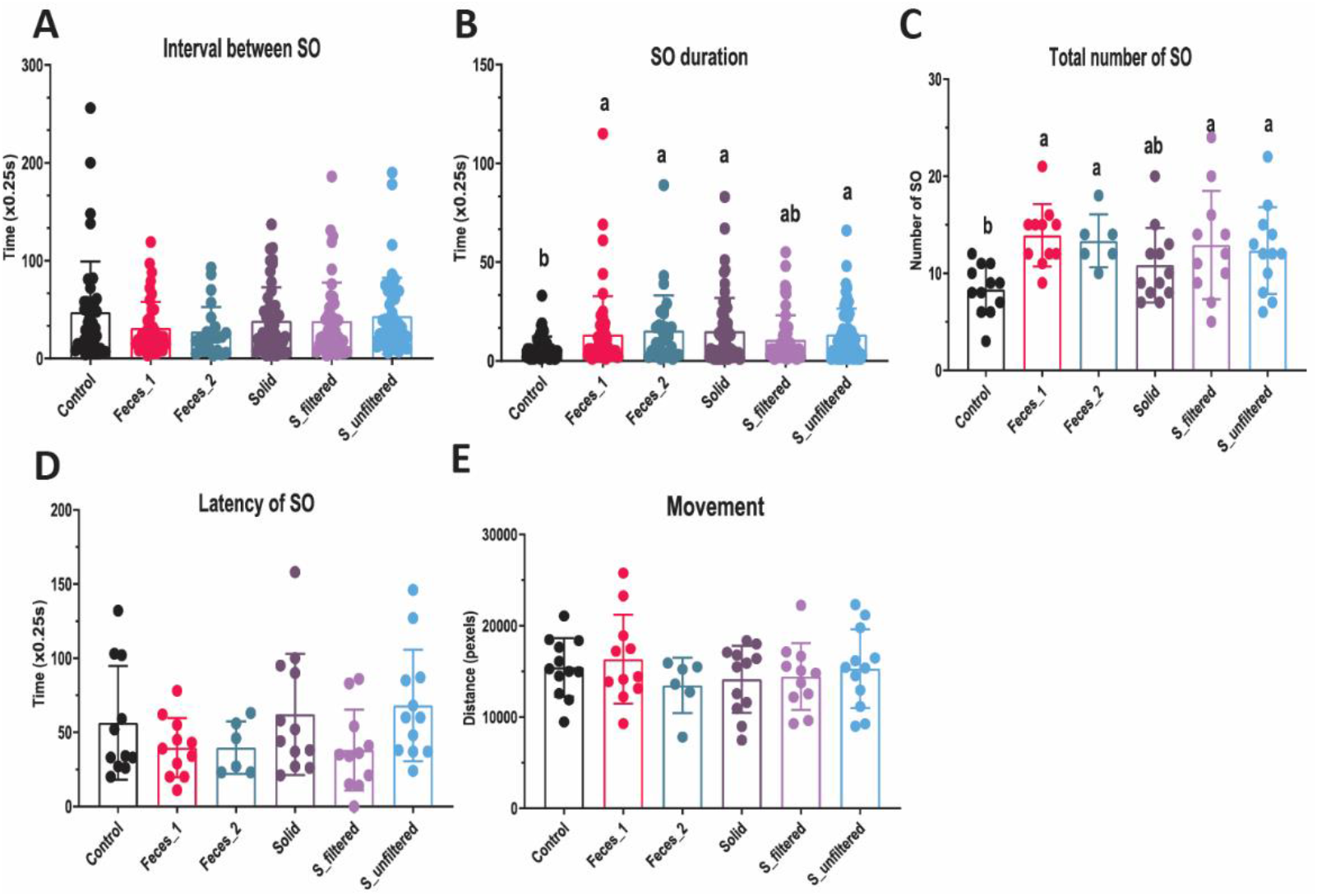
Additional analysis of male mouse behavioral responses to snake feces and its extracts. **A**, Interval between two consecutive SOs. **B**, Duration of each SO. **C**, Total number of SO. **D**, Latency of the first SO. **E**, Total movement distance. * indicates *p* < 0.05. For this experiment, 6 mice were available for analysis.

## Discussion

Recent advances in ML have tremendously facilitated our ability to measure and understand animal behavior (Wang, 2019). Multiple ML-based software packages, such as DeepLabCut (Mathis et al., 2018a), Social LEAP (SLEAP) (Pereira et al., 2022), and DeepPoseKit (Graving et al., 2019), can efficiently and accurately extract animal movement and posture information from videos. Despite this progress, there are still challenges for analyzing and interpreting complex, diverse, and high-dimensional behavior data sets (Pérez-Escudero et al., 2014; Nakamura et al., 2016). The layered, hybrid analytic workflow presented here was developed with the goal of incorporating the advantages of supervised and unsupervised ML classifiers to improve the accuracy and interpretability of animal behavior quantification.

In general, the objective of a study and the format of behavioral data determine the best analytic strategies and tools. In many cases, experimenters carefully design conditions that they hypothesize will generate changes in pre-defined behavioral states of interest. In these cases, supervised classifiers, such as RF, are generally well-matched to the overall goal (Nilsson et al., 2020; Winters et al., 2022). In our study, a well-trained RF achieved high initial accuracy (>95%), but had high error rates in highly dynamic/transient states (e.g. LV and AP). Because RF classifiers do not integrate temporal/sequential components in their predictions, we applied a series of rolling median filters to input data to create temporally smoothed copies of the animal’s body positions and orientations (i.e. some temporal information). This boosted the classification performance (to 0.9931) and improved prediction accuracy for transient states (e.g. LV and AP). Since animal movement is highly dynamic, and often includes fast-switching between behavioral states (Fauchald and Tveraa, 2006), improving the accuracy of supervised models in classifying transient states has great value.

In experiments where the experimental goal is to more generally explore the structure of animal behavior, unsupervised models, such as HMMs, are well-suited (Wiltschko et al., 2020; Findley et al., 2021). The main caveat of HMMs is that the outcomes may not map directly to specific behaviors of interest. In this study, our experimental design was specifically intended to assess mouse threat assessment, but HMM states mapped poorly onto our user-defined ethogram (GT states; Fig. 3). Increasing the number of HMM states, in hopes of finding some latent states with high interpretability, did not succeed (Fig. 3). This experience was the primary motivation for pursuing a hybrid model. We found that the reHMM model improved interpretability, even with a small number of states (Fig. 4A-B). reHMM+, which incorporated RF, HMM, and a single high-importance input feature, resulted in excellent interpretability, nearly matching user-defined GT states (Fig. 4C-D). The high degree of interpretability of reHMM+ comes with the added benefit of being backed by a dynamic model. The layered architecture reduced burdens (computational time and end-user evaluation) associated with determining the best parameters for HMM training and application (e.g. the number of hidden states). Furthermore, by taking advantage of high-value but low-computational cost components (RF output, high-importance raw features), reHMM+ has substantially reduced complexity and computational cost, making it highly flexible and adaptive.

Our behavioral tests demonstrated that reHMM+ is well-suited for behavioral pattern analysis in mouse risk assessment behavior. TMT-induced behavioral responses in these conditions ran counter to typical findings, specifically that TMT causes hiding/freezing and total odor avoidance (Morrow et al., 2000). In our conditions, TMT-exposed mice tended to stay away from the odorant, but adopted a rapid-and-short investigative pattern to sample (SO) TMT (Fig. 5). Using reHMM+, we also find that mice react to potentially risky cues (snake feces and its extracts) with a different pattern of sampling and exploring than TMT, consistent with risk assessment (Fig. 6). A major benefit of reHMM+ in this context is that traditional measurements from user-defined ethograms (time spent in state, number of times entering state) can be evaluated along with state transition information, producing a broad and deep behavioral profile. Combined, the data generated by the reHMM+ workflow are capable of distinguishing responses to odors with similar overall valence but variably overlapping molecular components (e.g. TMT versus snake feces, etc.). In the future, we anticipate that this analytical approach will allow identification of novel behavioral effects of chemosensory secretions, and ultimately the brain processes that underlie diverse chemosignal-mediated behaviors.

The benefits of reHMM and reHMM+ workflows make them attractive in experiments designed to investigate specific behavioral hypotheses, but they also have several noteworthy limitations. First of all, we developed reHMM+ using specific ML models with complementary strengths and weaknesses (e.g., HMM and RF). Although reHMM and reHMM+ models do not require extensive tuning of model hyperparameters, their performance still varies depending on several factors, such as feature selection and the specific behavioral ethogram chosen. Additionally, the choice of the tracking feature to add into the reHMM+ layer may qualitatively affect the end result. Here, the reHMM+ model incorporates the most important feature in RF classification – the distance from the mouse nose to the sample dish. This choice resulted in HMM states that closely matched the GT (ethogram) states, improving interpretability. This had the consequence of seemingly reducing the influence of the first-order HMM states (Figs. 4-5). Since a major benefit of HMMs is objective assignment of latent states that may have undiscovered important neurobiological underpinnings (Leos-Barajas et al., 2017), the reHMM+ output may not always be superior to reHMM. The value of the reHMM+ approach depends on the degree to which investigators wish to adhere to subjective, but interpretable, behavioral states.

Overall, we found that applying layered, hybrid ML workflows in our experimental context unveiled distinctive mouse behavioral patterns induced by established and experimental chemosensory stimuli, indicating diversity in the way chemical signals modulate mouse risk assessment behavior. Because of their inclusion of user-specified ethograms, we anticipate that the reHMM and reHMM+ models will be especially powerful in the context of more complicated experimental settings, such as multi-animal social interaction tests. Ultimately, we find that layered, hybrid analytic workflows improve the depth and reliability of ML classifiers and expand the ability to explore the dynamic architecture of animal behavior.

## Acknowledgements

We thank the Dallas Zoo Department of Herpetology for providing reptile biological samples (DZ Project S2019-2). We thank members of the Meeks Laboratory and BioHPC for constructive feedback.

